# Psi represses EGFR signalling in the neural stem cell niche to inhibit neuroblast proliferation in the *Drosophila* brain

**DOI:** 10.64898/2026.04.01.715755

**Authors:** Damien Muckle, Brooke Kinsela, Tanya Javaid, Nan-hee Kim, Naomi Mitchell, Teresa Bonello, Leonie M. Quinn, Olga Zaytseva

## Abstract

In mammals, neural stem cells (NSCs) generate the neurons and glia essential to brain development, while defective NSCs drive brain cancer (glioma). NSC fate is not only controlled cell intrinsically, but also relies on signalling from the cellular microenvironment, or stem cell niche. Defective communication between NSCs and their niche can, therefore, drive stem cell renewal over differentiation to promote glioma. Here, we demonstrate that the orthologue of glioma-driver mutation FUBP1, *Drosophila* Psi, is essential in the cortex glia, the NSC niche, for preventing NSC overproliferation. We further demonstrate that Psi controls NSC fate through direct transcriptional repression of EGFR ligands, spitz (spi) and gurken (grk); with spi required cell intrinsically to enable proliferative growth of cortex glia, while grk functions cell extrinsically to control NSC fate. These observations highlight the complexity of EGFR function in modulating communication between NSCs and their niche. Our findings demonstrate the importance of understanding the nuances of intrinsic and extrinsic control of NSC fate, mechanisms of cell-cell communication critical to animal development that will provide insight into FUBP1/EGFR-driven glioma.

## Introduction

In the brain, neural stem cells (NSCs) self-renew to maintain the stem cell pool and differentiate to generate the neurons and glia comprising the central nervous system (Gage and Temple, 2013). The delicate balance between NSC proliferation, differentiation and quiescence is controlled by a combination of intrinsic genetic and epigenetic programs (Hsieh and Zhao, 2016; Renault et al., 2009). Furthermore, NSC fate is controlled by extrinsic cues from the surrounding cellular microenvironment, or stem cell niche (Li and Guo, 2021; Ma et al., 2005). Dysregulation of the endogenous paracrine signalling and structural support from the niche in the control of NSC fate likely contributes to initiation of primary brain cancers such as glioma (Liu & Zong, 2012; Ligon et al., 2007). Indeed, the low-grade glioma, oligodendroglioma, is predominantly comprised of differentiated glial-like cell populations, including astrocytes and oligodendrocytes, in addition to small populations of glioma stem and progenitor cells (Tirosh et al., 2016). Based on NSC fate control during normal development (Hsieh and Zhao, 2016; Li and Guo, 2021; Renault et al., 2009), tumour progression is likely driven by undifferentiated glioma stem cells, generated through combined disruption of intrinsic mechanisms and extrinsic cues from the surrounding glioma stem cell niche, predominantly comprised of glia.

Loss-of-function *FUBP1* mutations are detected in, and predicted to drive, oligodendroglioma (Bettegowda et al., 2011; Brat et al., 2015; Chakravarty et al., 2017; Chang et al., 2016; Liu et al., 2023). The *FUBP* family encodes dual function proteins, capable of binding single stranded DNA to regulate transcription, and RNA to control splicing and translation (Debaize and Troadec, 2019; Hoang et al., 2019; Jacob et al., 2014; Liu et al., 2011; Rabenhorst et al., 2009; Rabenhorst et al., 2015; Zheng and Miskimins, 2011; Zhou et al., 2016). FUBP1, the prototypical family member, functions as an oncogene in most cancers, at least in part, through direct transcriptional activation of *MYC* (Avigan et al., 1990; Duncan et al., 1994; Wang et al., 2021; Wang et al., 2022; Xiong et al., 2024; Zhang et al., 2023) and additional pro-proliferative targets, including *c-KIT* (Debaize et al., 2018) and Wnt/β-catenin pathway activators (Yin et al., 2021). Nevertheless, the mechanistic basis for FUBP1’s function as a tumour suppressor, rather than an oncogene, in oligodendroglioma, is poorly understood. Supporting *in vivo* functional evidence is limited to histological analysis of *Fubp1* knockout embryonic mouse brains, which display impaired development suggestive of hyperproliferation (Zhou et al., 2016), although the molecular mechanisms underlying hyperplasia in the *Fubp1*-null embryonic mouse brain remain unclear. Furthermore, *Fubp1* knockdown in mouse neural progenitors results in impaired differentiation through altered splicing of KDM1A/LSD1 (Hwang et al., 2018), although direct genome-wide regulatory targets of FUBP1 remain undefined.

FUBP1 is structurally and functionally conserved as a single orthologue in *Drosophila*, P-element somatic inhibitor (Psi) (Siebel et al., 1994) which, in line with FUBP1’s role in activating *MYC* transcription, directly binds and regulates *Myc* transcription to promote cell and tissue growth in the developing wing (Guo et al., 2016; Zaytseva et al., 2023). In addition to driving growth via Myc, genome-wide DNA binding and RNA sequencing studies revealed that Psi orchestrates cell and tissue growth in the developing wing through increased expression of the mitotic phosphatase *string*, and transcriptional repression of growth inhibitors, including *tolkin* (Zaytseva et al., 2023).

In the *Drosophila* larval brain, the niche is comprised of cortex glia, which forms an extensive network encasing neural stem and progenitor cells (NSPCs) to provide the structural support, signalling cues and protection from stress required for NSPC renewal and differentiation (Corty and Coutinho-Budd, 2023; Egger et al., 2008; Homem and Knoblich, 2012; Yuan et al., 2020). Here, we demonstrate tumour suppressor functions for Psi in the larval brain, whereby Psi is required cell intrinsically to promote proliferation in the cortex glia (i.e. NSC niche), yet functions cell extrinsically to prevent NSC overproliferation. These observations demonstrate that Psi functions in an oncogenic manner in cortex glia while, concurrently, impairing renewal of neighbouring NSCs i.e. functioning as a tumour suppressor. We further demonstrate that Psi represses EGFR ligands to not only control proliferative growth of cortex glia (spi) but to also regulate communication with NSCs (grk). Thus, Psi is normally required in the cortex glial niche to control NSC fate via the EGFR ligand grk; an observation with potential to provide clues into oligodendroglioma initiation and/or progression driven by FUBP1 loss-of-function. Furthermore, our data suggests FUBP1 loss-of-function drives glioma, at least in part, by dysregulating EGFR signalling between NSCs and their glial niche.

## Results

### Psi is required in the cortex glial niche to prevent neural stem cell expansion

Prior to investigating potential roles for Psi in the cortex glia, which comprise the NSC niche, we monitored Psi expression in the larval brain using our previously validated custom anti-Psi antibody (Guo et al., 2016; Zaytseva et al., 2023). We observed Psi expression in both cortex glia (white arrowhead) and NSCs (yellow arrowhead) of the third instar brain lobe (**Figure 1A, B**). Efficiency of Psi knockdown (KD) was demonstrated for two independent Psi RNAi lines at the mRNA level using qPCR (**Figure 1C**) and protein level depletion was observed using the anti-Psi antibody, following Psi KD of either RNAi line driven by *GMR54H02*-GAL4, i.e. *wrapper-*GAL4 (**Figure 1D**). *wrapper-*GAL4 driven Psi KD in cortex glia significantly reduced glial cell number for both Psi RNAi lines (**Figure 2A, B**). In contrast to the decrease in glia, quantification of NSCs revealed Psi KD in the cortex glial niche significantly increased abundance of neighbouring NSCs (**Figure 2C**) and, moreover, the proportion of NSCs in mitosis (**Figure 2D**). These data suggest Psi is required cell intrinsically for promoting proliferative growth of cortex glia, while function in the niche is essential for limiting renewal of neighbouring NSCs.

**Figure 1.**
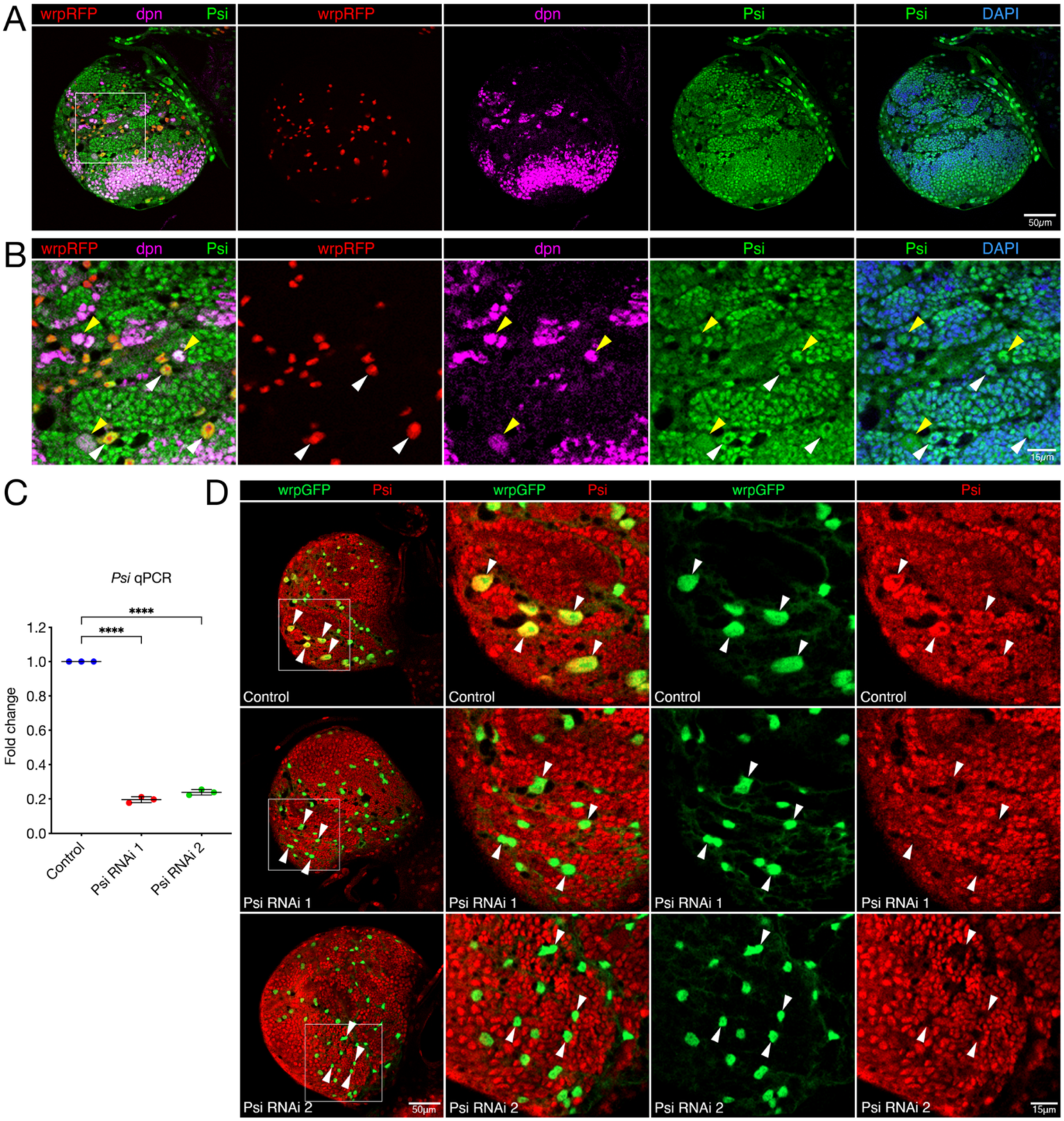
Psi is expressed in NSCs and glia in the third instar larval brain. (A) Third instar larval brain with cortex glia labelled by *wrapper*-GAL4-driven *UAS-mCherry.NLS* (wrpRFP) and NSCs marked by deadpan-GFP protein trap (dpn, false coloured to magenta). Psi protein is stained with anti-Psi polyclonal antibody (green) and DNA is stained with DAPI (blue). (B) Inset from white box in 1A. White arrowheads indicate cortex glia and yellow arrowheads indicate NSCs. (C) qPCR for *Psi* mRNA in third instar larval brains after ubiquitous expression of alternate and non-overlapping *UAS-Psi* RNAi lines: *UAS-Psi* RNAi 1 (V105135) or *UAS-Psi* RNAi 2 (V28990) driven by *tubulin*-GAL4 after inactivation of GAL80^ts^ for 48 hours compared with control. Relative mRNA levels normalised to the average of *Cyp1* and *tubulin* reference genes (n=3, ****p<0.0001). (D) Psi protein for control or following cortex glial-specific Psi depletion using *UAS*-*Psi* RNAi 1 or *UAS*-*Psi* RNAi 2 by *wrapper*-GAL4. Cortex glia labelled with *wrapper-*GAL4-driven *UAS-GFP.NLS* (wrpGFP, white arrowheads) and Psi protein stained using anti-Psi polyclonal antibody (red).

**Figure 2.**
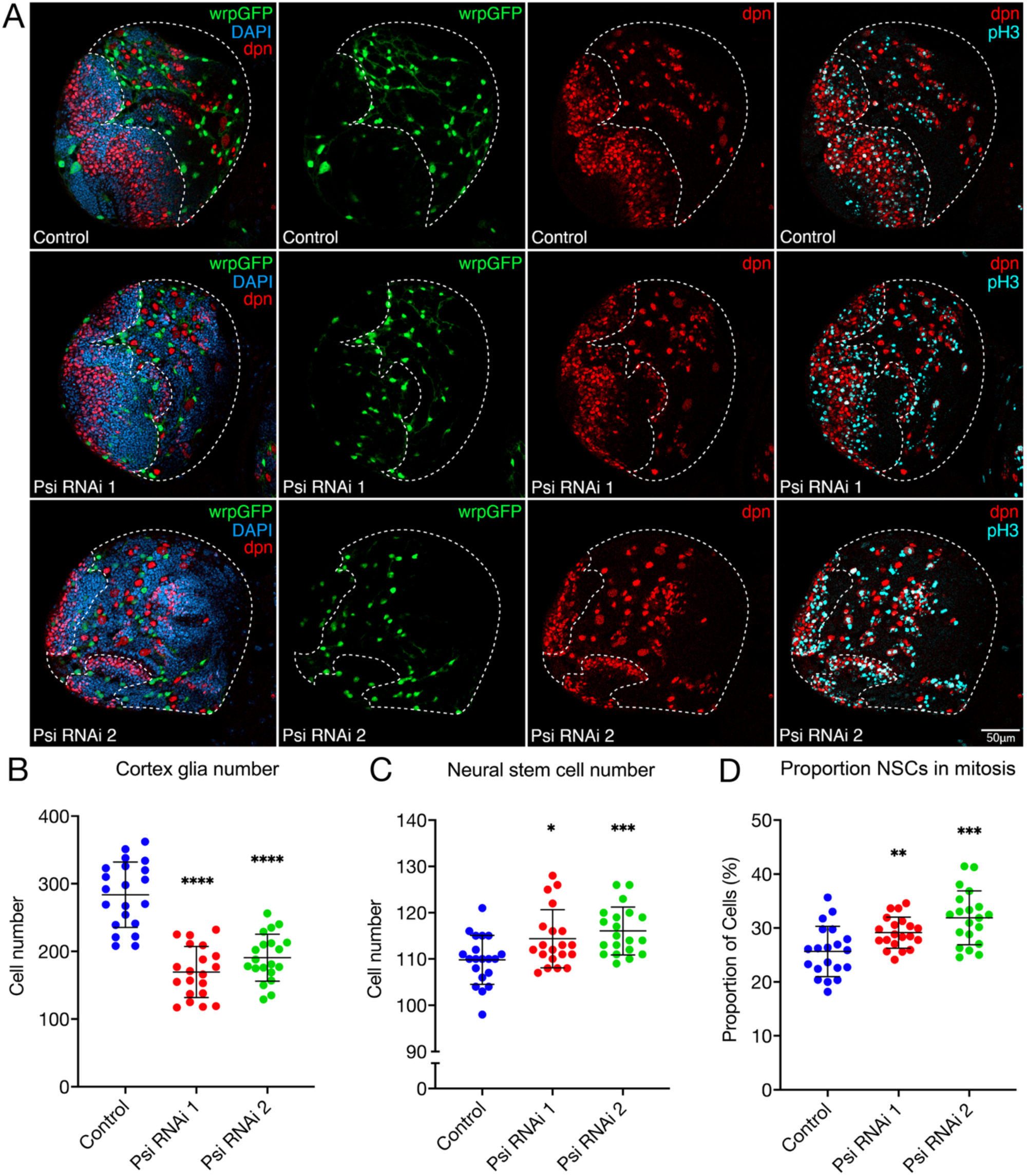
Psi knockdown cell intrinsically reduces cortex glial growth and cell extrinsically drives expansion of NSCs. (A) Third instar central brains (outlined) with *wrapper-*GAL4-driven Psi RNAi 1 or Psi RNAi 2 compared with control. Cortex glia (wrpGFP, green), NSCs (dpn, red), mitoses (pH3, cyan), DNA (DAPI, blue). Scale bar = 50 µm. For central brains with *wrapper-*GAL4-driven Psi RNAi 1 or Psi RNAi 2 compared with control: (B) cortex glia number, (C) NSC number and (D) proportion of NSCs in mitosis (n≥20, ****p<0.0001, ***p<0.001, **p<0.01, *p<0.05, Student t-test, error bars represent SD).

### Psi KD alters expression of cortex glial cell growth, migration, and signal transduction genes

To determine the Psi KD transcriptome underpinning cell intrinsic function in glial growth and extrinsic control of NSC fate, we used bulk RNA sequencing (RNA-seq) analysis of FACS-isolated Psi KD, compared with control cortex glia (**Figure 3A**, see **Figure S1** for QC). DESeq2 analysis (Love et al., 2014) revealed differentially expressed (DE) genes, which were both up- and down-regulated (53% and 47%, respectively) in Psi KD cortex glia (**Figure 3B-C**, FDR<0.01, **Data S1**). Consistent with Psi KD reducing cortex glial cell number (**Figure 2**), genes upregulated by Psi KD included regulators of cell growth and metabolism (**Figure 3D)**. Moreover, DE genes were enriched for pathways implicated in cell migration, actin cytoskeleton organisation, morphogenesis, and signal transduction (**Figure 3D-E, Data S1**), consistent with Psi functioning in cortex glia to control communication with NSCs and the observed increase in NSCs associated with Psi KD (**Figure 2**).

**Figure 3.**
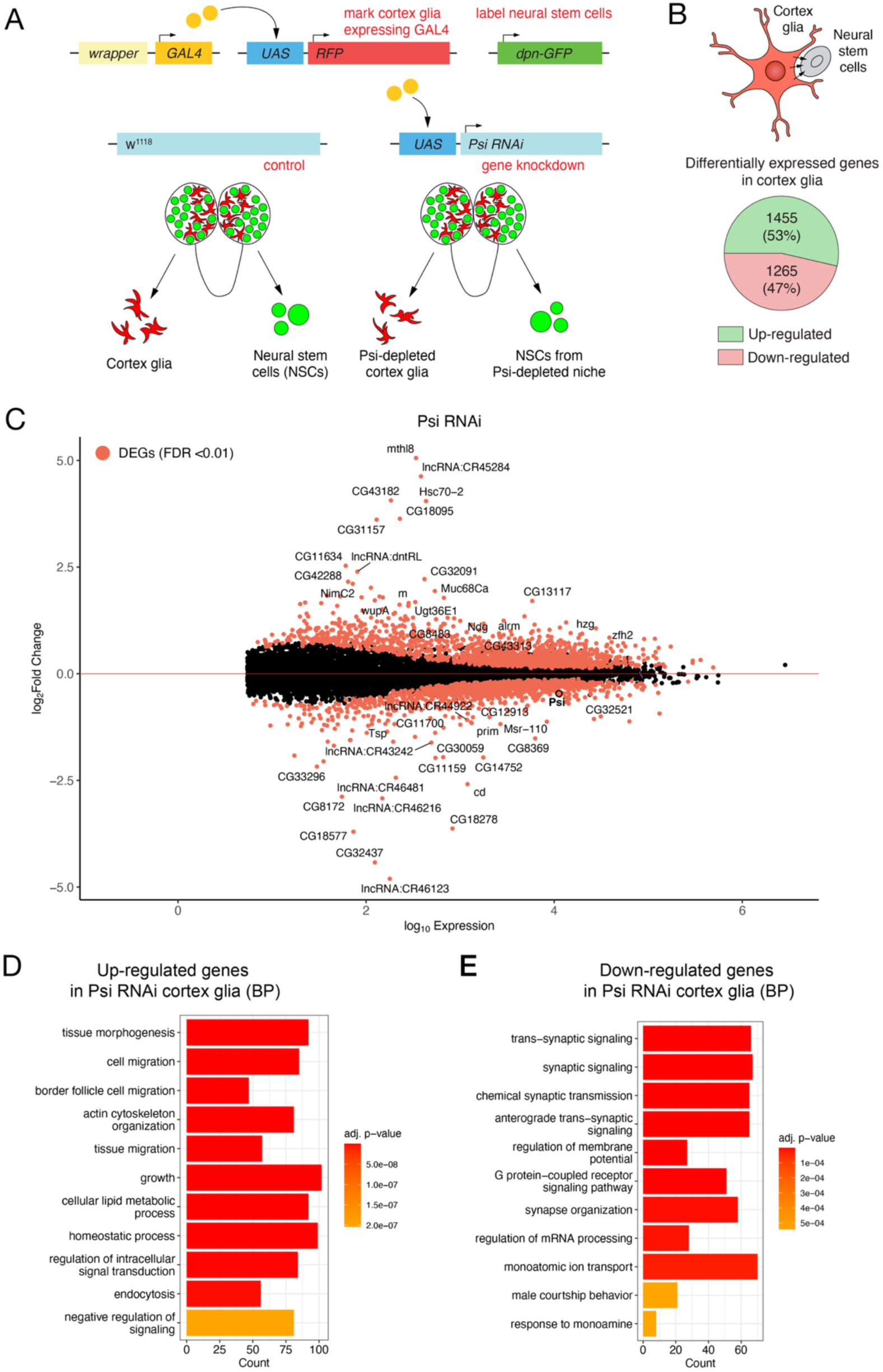
Genes differentially expressed in Psi KD cortex glia. (A) Transcriptome profiling of cortex glia and associated NSCs. (C) Total number of genes upregulated or downregulated in cortex glia. (D) MA plot for all differentially expressed genes in Psi KD cortex glia. Significantly differentially expressed (DE) genes (FDR<0.01) in red and the 50 genes with the greatest fold change are labelled. (D) Biological process (BP) ontology of DE genes upregulated and (E) downregulated in Psi KD cortex glia.

Additionally, in line with Psi’s established function in RNA splicing control (Adams et al., 1997; Labourier et al., 2002; Siebel et al., 1994; Siebel et al., 1995; Wang et al., 2016), 858 genes were differentially spliced in Psi KD cortex glia, with exon skipping the most common alteration (**Figure S2A, Data S2**). Differentially spliced genes were implicated in cell growth, regulating cell junctions, and cell-cell signalling (e.g. including cell-adhesion protein armadillo and cell polarity kinase discs large) (**Figure S2B, Data S2)**. Nevertheless, intersection of differentially expressed Psi targets and alternatively spliced genes revealed that the majority (88%) of transcriptional changes occur independently from splicing (**Figure S2C**), consistent with the predicted dual roles for Psi (i.e. transcription and splicing) in gene regulation in glia.

### Direct transcriptional targets of Psi are implicated in cell growth, migration, and signalling

To identify Psi’s direct transcriptional targets, we profiled Psi’s genome-wide binding targets specifically in cortex glia using Targeted DamID (TaDa) (Marshall et al., 2016; Southall et al., 2013) (**Figure 4A**), for intersection with the DE genes associated with Psi KD in cortex glia identified by RNA-seq (**Figure 4B**). To identify Psi’s cortex glial targets, the Dam-Psi fusion protein was driven with *wrapper*-GAL4, and DNA binding determined using established peak calling methodologies (Marshall and Brand, 2015; Marshall et al., 2016). After normalising to the Dam alone control and confirming low variability between replicates (>0.8 Spearman correlation, **Figure S3A**), we identified 1848 genes bound by Psi with statistical significance (**Figure 4C**, **Data S3**). Ontology analysis of Psi’s direct targets in cortex glia revealed enrichment for genes implicated in development, morphogenesis, cell migration, and actin cytoskeleton organisation (**Figure S3B, Data S3**).

**Figure 4.**
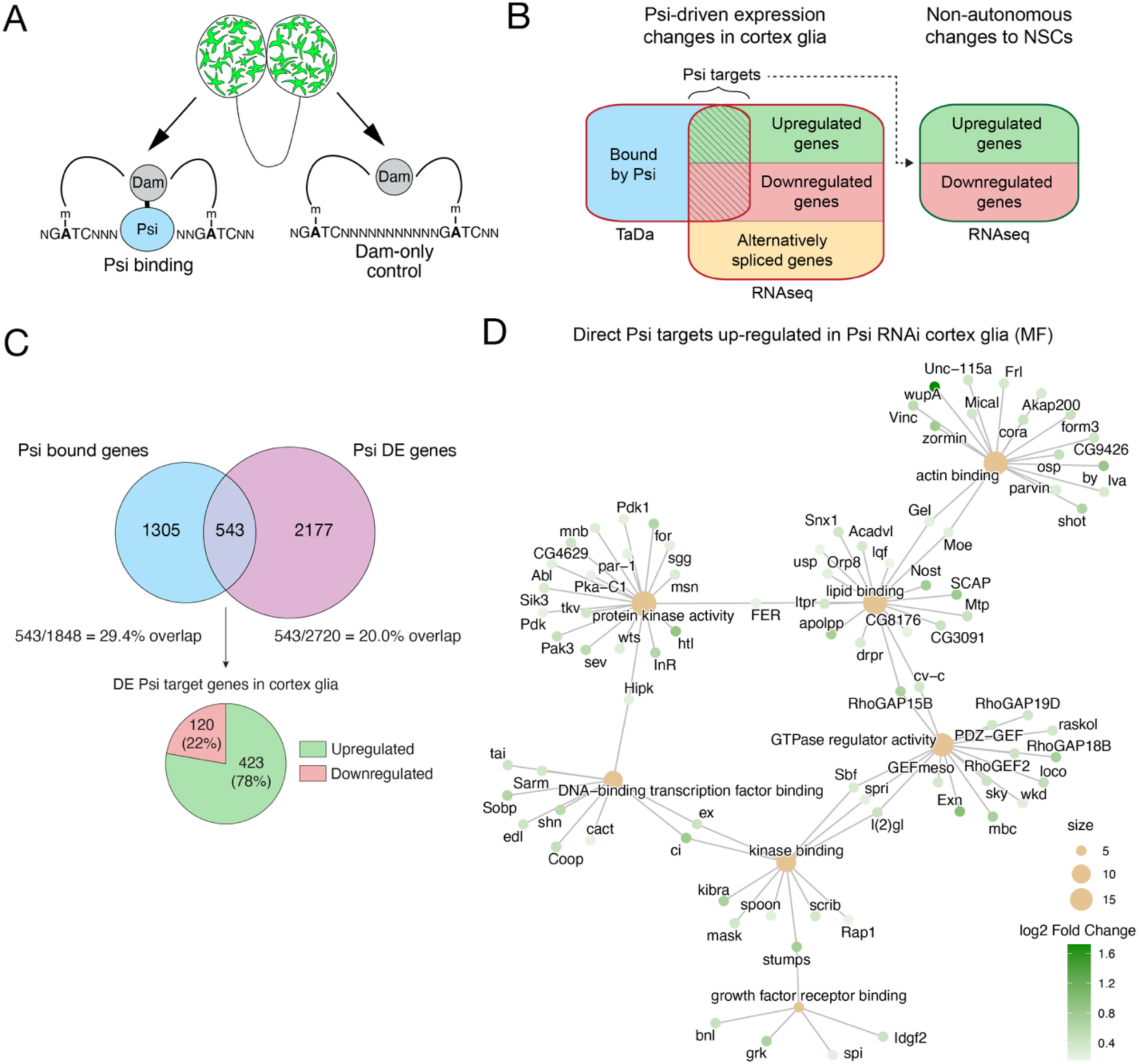
Psi directly binds genes involved in cell communication in cortex glia. (A) TaDa profiling of Psi genome-wide in cortex glia. (B) Workflow for identification of direct Psi targets in cortex glia and indirect expression changes in NSCs. (C) Differentially expressed (DE) direct targets of Psi in cortex glia identified by intersection of genes bound by Psi and genes DE following Psi KD in cortex glia. (D) Enriched molecular function ontology classes and the associated genes among direct Psi targets upregulated following Psi KD in cortex glia.

To identify direct Psi targets, differentially expressed (DE) by Psi KD, Psi bound genes were intersected with DE genes. Of the 543 genes shared between datasets (**Figure 4C, Data S4)**, most direct DE targets were upregulated by Psi KD (423 genes or 78%, **Figure 4C**), suggesting that Psi principally functions as a transcriptional repressor in cortex glia. Of note, most DE genes (2177 genes, 80%) were not bound by Psi and, thus, reflect changes in gene expression, or RNA stability, resulting from indirect effects of Psi depletion. Conversely, 70.6% of Psi bound genes were not differentially expressed (1305 genes), likely, at least in part, because Psi-Dam DNA interactions measured over a 24-hour time frame detects transient binding, which will not always be associated with significantly altered gene expression.

Gene Ontology (GO) analysis revealed direct Psi targets, upregulated in Psi KD cortex glia, were enriched for the biological processes of cell projection, morphogenesis, adhesion and cell migration, (**Figure S4A, Data S4)**; observations consistent with cell extrinsic roles for Psi in the niche for controlling renewal of neighbouring NSCs (**Figure 2**). Although significant gene set enrichment was not observed for Psi’s direct downregulated targets, upregulated genes included gene groups implicated in growth factor receptor binding, including genes encoding EGFR ligands, spi and grk (**Figure 4D, Figure S4B**). Furthermore, in accordance with the observed expansion of neighbouring stem cells, and identification of gene clusters implicated in EGFR/growth factor receptor binding, KEGG pathway enrichment analysis revealed upregulation of multiple EGFR/MAPK pathway components in Psi KD cortex glia (**Figure S5**).

### Psi KD in cortex glia cell non-autonomously alters the NSC transcriptome

In addition to analysis of cell intrinsic changes to Psi KD cortex glia, transcriptional profiling was performed on FACS isolated dpn-GFP positive cells (i.e., type I and II NSCs and mature INPs of the type II lineage) (**Figure 3A**). RNA-seq of these neural stem and progenitor cells (NSPCs) associated with the Psi KD niche, compared with control, identified significant differential expression of 1378 genes (**Figures 5A, B, Data S5**), which were approximately equally up- or downregulated (**Figure 5A**). Ontology analysis of gene sets differentially expressed in NSPCs identified enrichment for signalling pathways, e.g. upregulation of EGFR pathway growth regulators *S6kII*, *Pi3K68D* and *phyl* (**Figure 5C, Data S5**), consistent with the NSC expansion associated with Psi KD in cortex glia (**Figure 2**).

**Figure 5.**
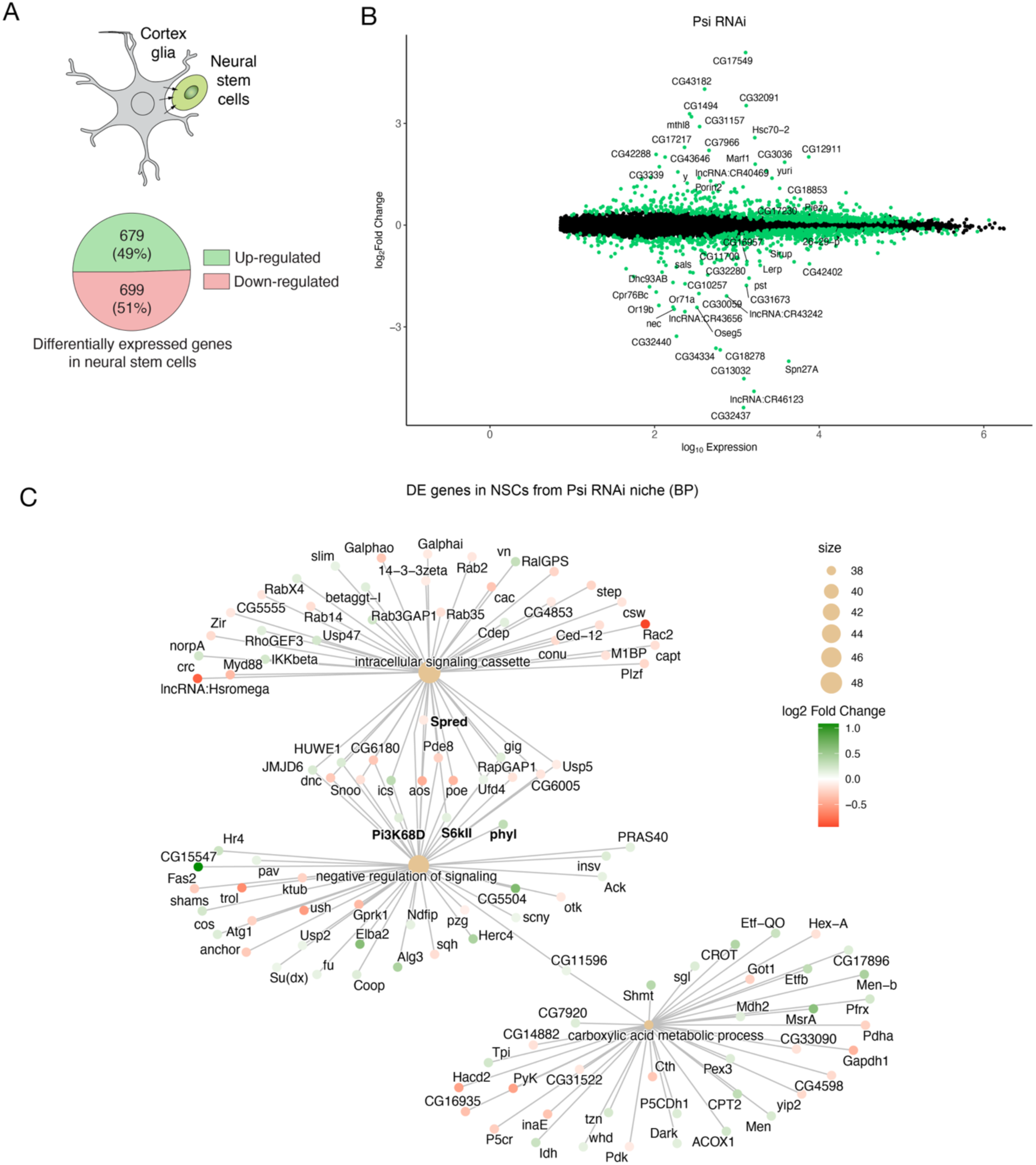
DE genes in NSCs associated with Psi KD cortex glia. (A) Upregulated vs downregulated genes in NSPCs associated with Psi KD in cortex glia. (B) Statistically significant DE genes in NSPCs (FDR<0.01) shown in green. The 50 genes with the greatest fold-change are labelled. (C) Enriched biological processes (BP) and DE genes in NSPCs associated with Psi KD cortex glia.

Moreover, in accordance with upregulation of EGFR ligands (e.g. spi and grk) in Psi KD cortex glia, which would be predicted to result in upregulation of ERK/MAPK signalling in neighbouring NSPCs, negative regulators of the ERK cascade were downregulated (**Figure 5C**). For example, we observed downregulation of *Spred*, whose mammalian orthologue inhibits growth factor-mediated activation of MAPK in mice (Wakioka et al., 2001), which would be predicted to activate MAPK and promote the NSC proliferation and expansion driven by cortex glial-specific Psi KD. Downregulated genes in NSCs were also enriched for neuron morphogenesis and differentiation (**Figure S6**), which would be expected to promote NSC renewal over differentiation. Together, these data suggest Psi regulates NSPC fate cell extrinsically by functioning in the cortex glial niche to provide both cellular support and signalling cues to control NSC fate.

### Psi repression of EGFR ligands, spi and grk, controls cortex glial and NSC fate, respectively

Next, we used functional genetic analysis to determine whether EGFR ligands, spi and grk, directly regulated by Psi and upregulated in Psi KD cortex glia, mediate the Psi KD cortex glial and/or NSC phenotypes. Previous studies have demonstrated that, in response to spi or grk, EGFR activates downstream signalling pathways (e.g. PI3K or MAPK), to control cell growth, proliferation and survival (Wee and Wang, 2017) (**Figure 6A**). Prior to testing whether the observed increase in spi mediates the impaired cortex glial proliferation and/or NSC expansion associated with Psi KD, we first confirmed that spi was efficiently knocked down via RNAi (**Figure S7A**). We then demonstrated spi KD in cortex glia significantly reduced cell number alone and, moreover, enhanced the Psi KD phenotype (**Figure 6B, C**). Spi KD in cortex glia did not, however, alter NSC number nor mitosis alone, nor did spi co-KD modify the NSC expansion phenotype driven by Psi KD (**Figure 6D, E**). Together these observations suggest spi ligand, produced in and secreted by cortex glia, normally functions in an autocrine manner to promote glial lineage proliferation.

**Figure 6.**
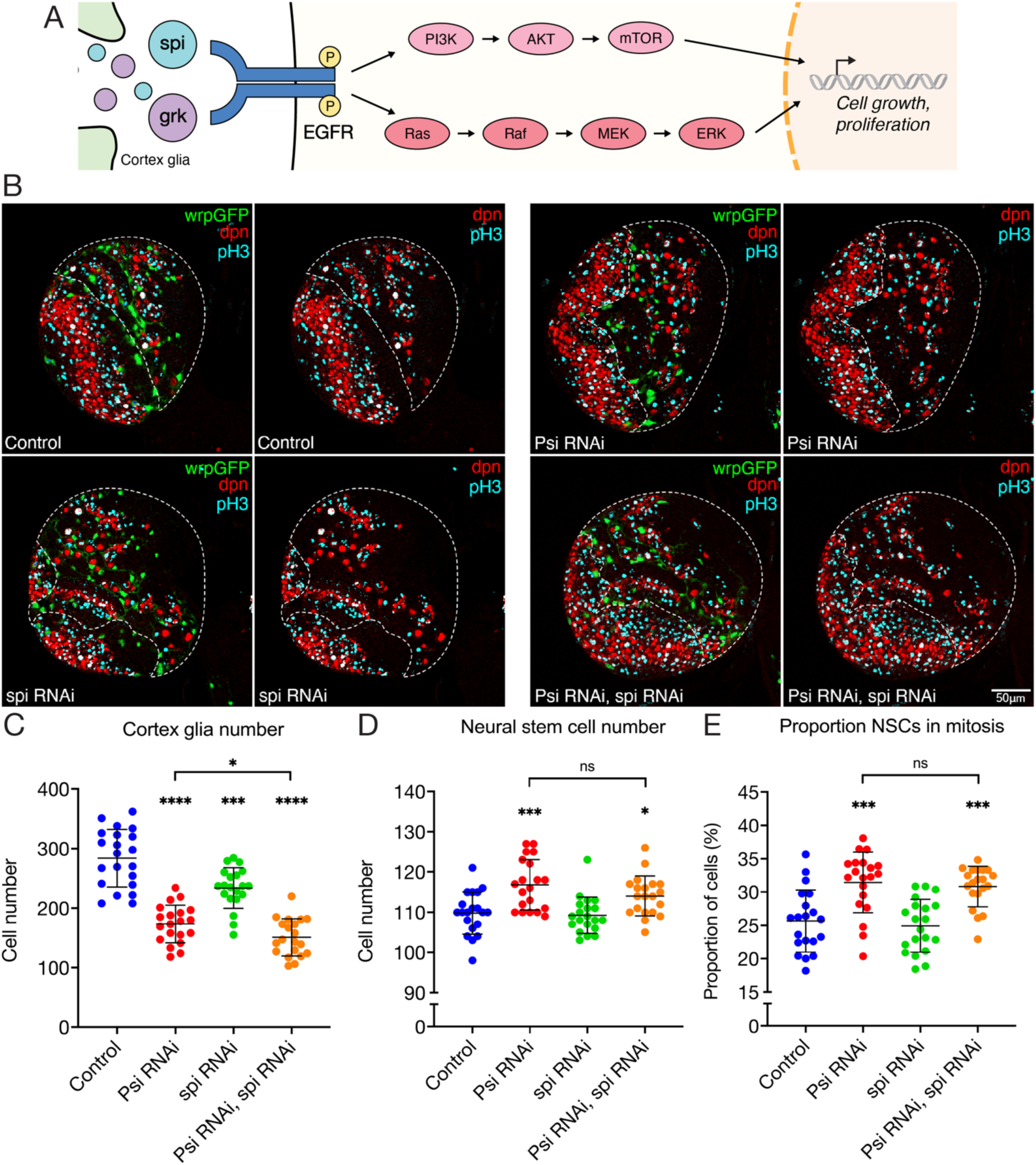
spi KD enhances the Psi KD cortex glial growth phenotype. (A) Schematic hypothesis: upregulated spi and grk in Psi-depleted cortex glia activate EGFR signalling. (B) Third instar central brains (outlined) with *wrapper-*GAL4-driven Psi RNAi, spi RNAi or co-KD compared with control. Cortex glia (wrpGFP, green), NSCs (dpn, red), mitoses (pH3, cyan), DNA (DAPI, blue). Scale bar = 50 µm. For central brains with *wrapper-*GAL4-driven Psi RNAi 1, spi RNAi or co-KD compared with control: (C) cortex glia number, (D) NSC number and (E) proportion of NSCs in mitosis (n≥20, ****p<0.0001, ***p<0.001, *p<0.05, ns=not significant, Student t-test, error bars represent SD).

Next, we tested whether grk RNAi, driven by *wrapper*-GAL4, altered cortex glial or NSC phenotypes alone or in combination with Psi KD. We first confirmed KD efficiency for the *grk* RNAi line (validated in **Figure S7B**) and then demonstrated grk KD in cortex glia did not alter cell number alone, nor did co-KD modify the reduced glial number associated with Psi KD (**Figure 7A, B**). Although grk KD in cortex glia alone did not alter NSC number, co-KD suppressed Psi KD-driven overproliferation and expansion of NSCs (**Figures 7A, C, D**). Therefore, in contrast with spi, the upregulation of grk associated with Psi KD in the niche was required for NSC expansion. Taken together, these data suggest the EGFR ligands spi and grk, which are directly repressed by Psi, have distinct roles downstream of Psi in the cortex glial niche: spi functions in an autocrine manner to mediate cortex glial proliferation, while grk ligands secreted from the niche function in a paracrine manner to mediate proliferation and expansion of neighbouring NSCs driven by Psi KD (**Figure 7E**).

**Figure 7.**
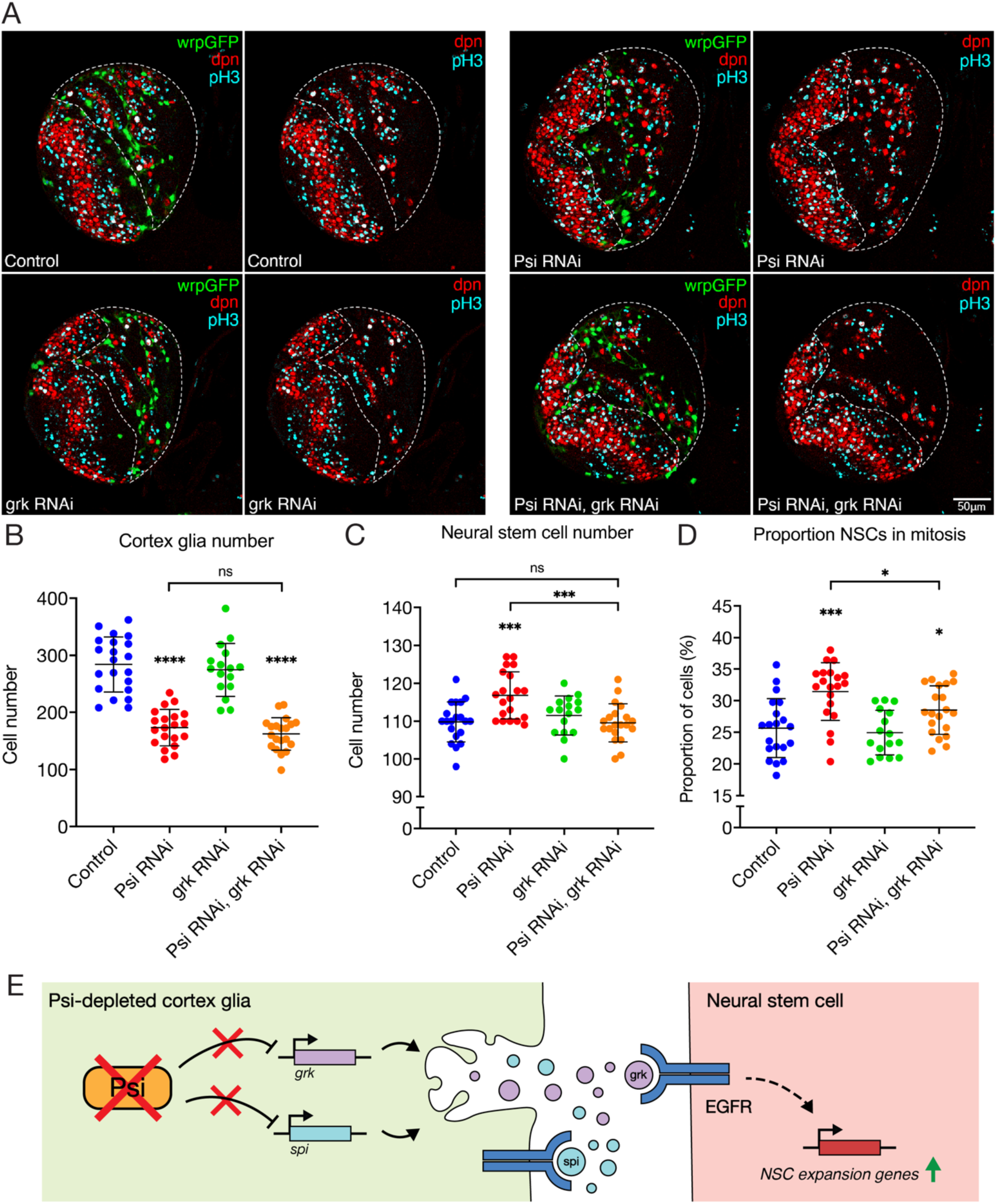
grk is required for NSC proliferation driven by Psi KD in cortex glia. (A) Third instar central brains (outlined) with *wrapper-*GAL4-driven Psi RNAi, grk RNAi or co-KD compared with control. Cortex glia (wrpGFP, green), NSCs (dpn, red), mitoses (pH3, cyan), DNA (DAPI, blue). Scale bar = 50 µm. For central brains with *wrapper-*GAL4-driven Psi RNAi 1, grk RNAi or co-KD compared with control: (B) cortex glia number, (C) NSC number and (D) proportion of NSCs in mitosis (n≥20, ****p<0.0001, ***p<0.001, *p<0.05, ns=not significant, Student t-test, error bars represent SD). (E) Model for spi and grk driving cell-intrinsic and cell-extrinsic proliferation in Psi-depleted cortex glia. Psi KD de-represses spi and grk. Cell-intrinsically, spi is required for growth and proliferation, while grk is required for growth. Subsequent grk upregulation in the Psi-depleted niche also drives increased cell proliferation of adjacent NSCs.

## Discussion

Although originally characterised as an oncogene (Zhang and Chen, 2012), FUBP1 functions as a tumour suppressor in oligodendroglioma: the first *FUBP1* allele is lost due to the characteristic whole-arm translocation t(1;19)(q10;p10), resulting in 1p/19q co-deletion, while the remaining allele is inactivated by subsequent somatic loss-of-function mutation (Bettegowda et al., 2011; Brat et al., 2015; Chakravarty et al., 2017; Chang et al., 2016; Liu et al., 2023). In accordance with FUBP1’s oncogenic functions, depletion of the *Drosophila* homologue Psi in cortex glia results in reduced cortex glia cell number. These data demonstrate Psi is required for proliferative growth of cortex glia in the developing brain, in accordance with the pro-proliferative function of Psi in the wing imaginal disc (Guo et al., 2016; Zaytseva et al., 2023). Of note, direct, differentially expressed targets associated with Psi-dependent growth control in the wing (e.g. *Myc*, *stg*, *dlp* and *tok*) (Zaytseva et al., 2023) were not among Psi’s direct differentially expressed targets in cortex glia. Therefore, despite Psi controlling growth in both contexts, this appears to be mediated by distinct growth-regulatory target genes.

The observation that spi KD alone was sufficient to significantly reduced glial cortex cell number, together with spi co-KD enhancing the Psi KD glial phenotype, suggests spi secreted by the cortex glia normally functions in an autocrine manner to control glial growth both downstream of, and in parallel to, Psi. In contrast, while optic-lobe cortex glia also secrete spi, in this context spi functions in a paracrine manner to activate EGFR signalling in the neighbouring neuroepithelium, driving expansion and promoting differentiation into neuroblasts (Morante et al., 2013). Thus, spi functions as an autocrine signalling factor in cortex glia in the central brain to control glial growth, while paracrine signalling from optic lobe cortex glia promotes expansion of the neuroepithelial stem cell pool.

The grk ligand was discovered, and is best characterised for, function as a paracrine signalling factor, secreted by germline cells to activate EGFR on neighbouring follicle cells in the ovary (Neuman-Silberberg and Schüpbach, 1993). Here, we demonstrate Psi directly represses transcription of the gene encoding the EGFR ligand grk in cortex glia, with the grk upregulation driven by Psi KD in the niche signalling in a paracrine manner to promote NSC expansion. Moreover, in accordance with the predicted upregulation of ERK/MAPK signalling in NSC associated with the Psi KD niche, we observed increased expression of downstream regulators of the EGFR pathway (*S6kII*, *Pi3K68D* and *phyl*) and downregulation of negative regulators of the ERK cascade, including *Spred*, whose mammalian orthologue inhibits growth factor-mediated activation of MAPK (Wakioka et al., 2001). These findings are consistent with Psi KD in the cortex glial niche activating EGFR/MAPK to promote the observed NSC proliferation and expansion.

Activated EGFR promotes proliferation, survival and migration, and inhibits neuronal differentiation in murine NSCs (Ayuso-Sacido et al., 2010) and is also a common feature of glioma (Saadeh et al., 2017), with EGFR amplification occurring in 35% of oligodendrogliomas (Wang et al., 2024). Given the high level of conservation between Psi and FUBP1 (Guo et al., 2016; Zaytseva et al., 2023), we predict FUBP1 loss-of-function will drive oligodendroglioma, at least in part, as a result of de-repression of EGFR ligand transcription.

Our observations that cell-cell communication between glia and stem cells are intriguing in context with oligodendroglioma, which is predominantly comprised of glial-like cancer cell populations, including astrocytes and oligodendrocytes, that provide structural support and paracrine signalling to control glioma stem cell (GSC) (Ligon et al., 2007; Liu and Zong, 2012; Tirosh et al., 2016). Taken together, given the high degree of functional homology with Psi, our findings suggest FUBP1 loss-of-function drives oligodendroglioma, at least in part, by increasing EGFR ligands in glial-like cells to promote GSC renewal. Thus, we predict communication between the glial-like cells and glioma stem/progenitor cells in glioma will contribute tumour initiation and progression.

Previous studies in *Drosophila* demonstrate glioma-like stem cells remodel their niche (Prager et al., 2020). For example, Delta expression in cortex glia is required to activate Notch on NSCs to drive terminal neural differentiation (Sood et al., 2024), while expression of *neurexin-IV* in the NSC lineage promotes encasement of NSCs by cortex glia (Banach-Latapy et al., 2023). Thus, in future, it would be of interest to determine whether Psi KD in the NSC lineage similarly alters glial lineage fate.

## Materials and Methods

### Drosophila melanogaster *stocks*

*Drosophila melanogaster* stocks were maintained on a standard molasses and semolina *Drosophila* medium. Genetic crosses were raised at 25°C except when performed in the background of temperature-sensitive GAL80 (GAL80^ts^) expression, where they were initially raised at 18°C followed by a shift to 29°C. The following lines were obtained from the Bloomington *Drosophila* Stock Centre (BDSC): w^1118^ (BDSC #3605), *GMR54H02 (wrapper)*-GAL4 (BDSC #45784), *tubulin*-GAL4 (BDSC #5138), *tubulin*-GAL80^ts^ (BDSC #7019), *UAS-mCherry.NLS* (BDSC #38425), *UAS-GFP.NLS* (BDSC #4776), deadpan-GFP (BDSC #59755). The *UAS*-*Psi* RNAi 1 (V105135), *UAS*-*Psi* RNAi 2 (V28990), *UAS*-*spi* RNAi (V3920) and *UAS-grk* RNAi (V36251) lines were obtained from the Vienna *Drosophila* Resource Centre (VDRC). TaDaG-Dam and TaDaG-Psi-Dam were generated as described previously (Zaytseva et al., 2023).

### Larval brain immunofluorescence, microscopy and image analysis

Heads of late third instar larvae were dissected in 1× Phosphate Buffered Saline (PBS) and fixed for 20 min in 4% w/v paraformaldehyde (PFA) diluted with 0.1% v/v Triton X-100 in PBS (PBT). Heads were blocked for 1 hour at room temperature in 5 mg/mL bovine serum albumin (BSA) and incubated overnight at 4°C with primary antibody. Primary antibodies used for immunofluorescence were: rabbit anti-Psi (1:500, custom-made), rat anti-deadpan (1:500, Abcam ab195173) and mouse anti-phospho-Histone3 (pH3) (1:2000, Abcam ab14955). Larval heads were incubated with fluorophore-tagged secondary antibody dilution for 2 hours, followed by staining with DAPI solution for 30 min at room temperature. Secondary antibodies used were: anti-rabbit 647 (1:500, Jackson ImmunoResearch, 711-605-152), anti-rat Rhodamine Red-X (Jackson ImmunoResearch, 712-295-153) and anti-mouse 647 (1:500, Jackson ImmunoResearch, 715-605-151). Washes between incubations were performed with PBT. Samples were stored in 80% glycerol in PBS at 4°C.

Larval brains were mounted to a consistent thickness of 62.5 μm between glass slide and coverslip in 80% glycerol. Brain lobes were imaged at 40× magnification in overlapping 1.2 μm sections using the Zeiss LSM800 confocal microscope (Zen Blue software). Fluorophores were imaged using band-pass filters to remove cross-detection between channels. Images were processed and prepared using Fiji and Adobe Photoshop. Larval brain Z-series images were 3D reconstructed and analysed using Bitplane Imaris 9.3.1. A 3D model of the central brain region was generated using manual perimeter tracing and the Surfaces tool. Cortex glia nuclei were counted using the Spots tool according to *wrapper*-GAL4-driven GFP fluorescence intensity and estimated nucleus diameter. Central brain NSCs were distinguished from dpn-positive intermediate neural progenitors according to position and size, and were manually counted using the Spots tool. Mitotic NSCs were quantified by isolating the anti-pH3 channel within the previously identified NSCs and counting NSCs with co-localised anti-dpn and anti-pH3 fluorescence. The proportion of NSCs in mitosis was calculated by dividing the number of pH3-positive NSCs by the total number of NSCs.

### qPCR

RNA was isolated from equivalent numbers of third instar larval brains (10 per genotype) using the Promega ReliaPrep RNA Cell Miniprep System and eluted twice in 20 μL nuclease-free water. RNA purity and integrity were assessed using an automated electrophoresis system (2200 TapeStation, Agilent Technologies). 7 μL eluted RNA and both Oligo-dT and random hexamer primers were used for each cDNA synthesis (GoScript Reverse Transcription System kit, Promega). qPCR was performed using Fast SYBR Green Master Mix (Applied Biosystems) using the StepOnePlus Real-Time PCR System and Sequence Detection Systems in 96-well plates (Applied Biosystems, 95°C for 20 sec, 40 cycles of 90°C for 1 sec and 60°C for 20 sec). Amplicon specificity was verified by melt curve analysis. Average Ct values for at least two technical replicates were calculated for each amplicon. Target gene expression was normalised to the mean of *Cyp1* and *tubulin* alone, which were selected for having high expression and low sample-to-sample variability, as determined by RefFinder. Fold change was determined using the 2^-△△Ct^ method.

Primers used were as follows:

Psi: 5’ CGATGGCATCCCATTTGTTTGT 3’ and 5’ GGTGGTCAAGACTACTCGGC 3’, spi: 5’ ACGCCCAGGCCCAATATTAC 3’ and 5’ GATCGGCTATCTTCACCGCA 3’, grk: 5’ AACAATTGTCGCCGTCACAGA 3’ and 5’ TCGGAGAAGTAAGATTCGGCTC 3’, tubulin: 5’ TCAGACCTCGAAATCGTAGC 3’ and 5’ AGCCTGACCAACATGGATAGAG 3’, Cyp1: 5’ TCGGCAGCGGCATTTCAGAT 3’ and 5’ TGCACGCTGACGAAGCTAGG 3’.

### Targeted DamID

Embryos from *tubulin*-GAL80^ts^ *wrapper*-GAL4 and *UAS*-Dam-protein fusion crosses were collected over the course of <4 h lays at 25°C, after which embryos were placed at the repressive temperature of 18°C for 7 days until larvae reached the second instar larval stage. Larvae were then shifted to 29°C for 24 h. Third instar larval brains were collected into cold PBS, treated with Proteinase K in 50 µM EDTA at 56°C for 1-3 h and genomic DNA was extracted using a Zymo Quick-DNA kit (D4069). DNA was digested at sites of GATC methylation using DpnI at 37°C overnight and purified using a Machery-Nagel PCR purification kit (740609.50). Samples were eluted into 30 µL H_2_O, of which 15 µL was used for PCR enrichment and library preparation. PCR adaptors were ligated to methyl-specific digestion sites at 16°C for 2 h using T4 DNA ligase. To minimise signal from unlabelled sites, unmethylated GATC motifs were digested using DpnII at 37°C for 2 h. PCR using

MyTaq polymerase (Bioline BIO-21,113) was performed with 3 long extension cycles followed by 17 short extension cycles (Vogel et al., 2007). PCR products were purified with a Machery-Nagel PCR purification kit and adaptors were removed by A1wI digestion at 37°C overnight. DNA samples were sonicated in 100 µL volumes using a Covaris S2 sonicator at 10% DUTY, 140 W peak incident, 80 duration and 200 cycles per burst, achieving a 300 bp average fragment size. Samples were purified and a library prepared using Sera-Mag Speedbeads hydrophobic carboxyl magnetic beads (GE Healthcare, 65152105050250). DNA concentrations were measured using a Qubit DNA HS assay kit (Thermo Fisher, Q32854) and <500 ng DNA for each sample was used to generate a library. DNA sequence ends were repaired with T4 DNA polymerase, T4 polynucleotide kinase and Klenow Fragment at 30°C for 30 min. 3’ ends were adenylated using Klenow 3’ to 5’ exo-enzyme at 37°C for 30 min. Unique index adaptors were ligated to each sample using NEB Quick Ligase at 30°C for 10 min and adaptor dimers were removed by cleaning samples twice using Sera-Mag beads. DNA sequences were enriched by PCR using NEB Next Ultra II Q5 Master Mix (NEB M0544S), before final clean up using Sera-Mag beads. Successful ligation of adaptors and the absence of adaptor concatemers were verified using an Agilent Bioanalyser, and the final concentration was measured using Qubit. Libraries were pooled to achieve an equimolar concentration of each sample and an overall library concentration of 2 nM. Samples were sequenced using a HiSeq2500 (Illumina) with 50 bp single-end reads.

TaDa data was analysed using the damidseq_pipeline workflow (Marshall and Brand, 2015). The pipeline workflow was used to align reads to the FlyBase release dmel_r6.39 reference genome with Bowtie2, to identify GATC sites, and to calculate the normalised log_2_ ratio of the Dam-fusion protein profile and Dam alone. Using the bigwig files generated in the pipeline and the deepTools package (Ramírez et al., 2016), Spearman sample correlation and genomic coverage clustered metaplots were generated. Average binding across the genome between three Dam-Psi replicates was determined by calculating average enrichment at each region flanked by GATC sites. Binding profiles were visualised in bedgraph format using the Integrative Genome Viewer (IGV). Significant binding peaks were determined at 1% FDR using the find_peaks script, genes within 1 kb of the peaks were identified using the peaks2genes script (Marshall and Brand, 2015).

### FACS and RNA sequencing

Single cell suspension for Fluorescence-Activated Cell Sorting (FACS) purification and RNA sequencing was conducted using a protocol adapted from Harzer et al (2013). Late third instar larval brains from parental crosses of dpn-GFP; *wrapper*-GAL4, *UAS-mCherry.NLS* and *UAS-Psi* RNAi or w^1118^ were collected after 6 days of GAL4-induced knockdown. Brains from dpn-GFP only, *wrapper*-GAL4, *UAS-mCherry.NLS* only and either *UAS-Psi* RNAi or w^1118^ flies were also extracted for use as single colour or negative controls in downstream FACS. For each sample, approximately 180 brains were extracted into Schneider’s *Drosophila* medium (Thermo Fisher) with 1% Penicillin-Streptomycin (Thermo Fisher) on ice. Following three washes in PBS, a dissociation solution containing 425 µL Schneider’s *Drosophila* medium supplemented with 1% Penicillin-Streptomycin, 50 µL 10 mg/mL Papain (Sigma, P4762) and 25 µL 20 mg/mL Collagenase I (Sigma, C2674) per sample was prepared (final concentration 1 mg/mL) and added to each sample. Samples were incubated at 30°C for 30 min on a rotating platform mixer. Following removal of the dissociation solution, samples were homogenised by repeated aspiration and expulsion in PBS using a pipette. Cells were pelleted by centrifuging tubes for 5 min at 5000 G at 4°C and the supernatant removed. Cells were washed by this process twice more in PBS and once in Schneider’s *Drosophila* medium. 1000 µL Schneider’s *Drosophila* medium was added to each sample and the total volume passed through a 35 µm cell strainer (Corning) into a 5 mL tube for FACS. Cortex glia and neural stem cells (NSCs) were purified from the single cell suspension according to the gating strategy shown in **Figure S1**. Cells were sorted into LBA lysis buffer with 2% 1-Thioglycerol (Promega ReliaPrep RNA Cell Miniprep System) using the BD FACSAria Fusion, and stored at −20°C prior to RNA extraction.

RNA was extracted using the Promega ReliaPrep RNA Cell Miniprep System kit and eluted in 20 µL nuclease-free water. Sample RNA integrity was verified using an Agilent Tapestation and RNA concentration measured using a Qubit RNA HS assay kit (Thermo Fisher, Q32852). Library preparation was conducted by the Biomolecular Resource Facility, John Curtin School of Medical Research, Australian National University. RNA was prepared using the TruSeq Illumina protocol with Oligo-dT mRNA enrichment and strandedness preservation. Samples were sequenced using the Illumina NovaSeq6000 with 150 bp paired-end reads. On basis of poor RNA yield and integrity, and low sequencing depth, cortex glia control replicate #4 was excluded from subsequent analysis.

### Differential expression analysis

RNA sequencing reads were aligned to the FlyBase release dmel_r6.39 reference genome using STAR 2.7.8a. Aligned reads were counted using Subread featureCounts 2.0.2 (Liao et al., 2014). Differential gene expression was analysed using the DESeq2 R package (Love et al., 2014), and significant events were identified using 1% FDR threshold.

### Differential splicing analysis

Differential alternative splicing was determined using rMATS 4.1.2 (Shen et al., 2014) with bam RNA sequencing alignment files previously generated as part of differential expression analysis. JCEC methodology (analysis with with both junction counts and exon counts) was used. Significant differential splicing events were identified using FDR<0.05.

### Gene ontology and KEGG pathway analyses

Ontology of genes identified in TaDa and RNA sequencing datasets was analysed using the clusterProfiler R package (Yu et al., 2012). Significantly enriched gene ontology categories from these genes were identified using a q-value cut-off of 0.05, which was corrected using the Benjamini-Hochberg FDR method. The clusterProfiler filtering function was applied to remove redundant parent ontology classes where applicable. KEGG pathway enrichment analysis was conducted using the Pathview R package (Luo and Brouwer, 2013).

### Statistics

All statistical tests excluding those in TaDa and RNA sequencing analyses were conducted using GraphPad Prism 10 software. All qPCR data and cortex glia and NSC phenotype analyses were conducted using unpaired two-tailed Student’s t-test with 95% confidence interval and Welch’s correct where applicable. For all phenotypic quantification, each datapoint is representative of a single third instar larval brain lobe. Brains from ≥ 3 independently repeated experiments were collected and pooled to control for biological variability. In all figures, error bars represent standard deviation and significance is denoted according to the GraphPad classification system: * (p = 0.01–0.05), ** (p = 0.001–0.01), *** (p = 0.0001–0.001) and **** (p< 0.0001).

## Acknowledgements

The authors acknowledge support from Microscopy Australia at the Centre for Advanced Microscopy, Australian National University, a facility enabled by National Collaborative Research Infrastructure Strategy (NCRIS), and the Biomolecular Resource Facility supported by NCRIS and the Australian Cancer Research Foundation (ACRF).

## Data Availability

Raw sequencing data files are available via the Gene Expression Omnibus (GEO) using accession numbers GSE320523 (DamID) and GSE320524 (RNA-seq).

